# Three-dimensional Structural Interrelations between Cells, Extracellular Matrix and Mineral in Vertebrate Mineralization

**DOI:** 10.1101/803007

**Authors:** Zhaoyong Zou, Tengteng Tang, Elena Macías-Sánchez, Sanja Sviben, William J. Landis, Luca Bertinetti, Peter Fratzl

**Author notes:** equal contribution (Zhaoyong Zou, Tengteng Tang). corresponding author: Peter Fratzl, **Email:**. **Author Contributions** Experimental design (ZZ, LB, PF), Conduct of experiment (ZZ [Sample preparation, confocal microscopy, FIB-SEM, micro-CT], TT [FIB-SEM, micro-CT], EM-S [Sample preparation, TEM], SS [Sample preparation], LB [FIB-SEM]), Data analysis (ZZ, TT, EM-S, WJL, LB), Manuscript composition (ZZ, TT, EM-S, WJL, LB, PF), All authors discussed, read and revised the manuscript.

## Abstract

The spatial-temporal relationship between cells, extracellular matrices and mineral deposits is fundamental for an improved understanding mineralization mechanisms in vertebrate tissues. By utilizing focused ion beam-scanning electron microscopy with serial surface imaging, normally mineralizing avian tendons have been studied with nanometer resolution in three dimensions with volumes exceeding tens of microns in range. These parameters are necessary to yield fine ultrastructural details while encompassing tissue domains sufficient to provide a comprehensive overview of the interrelationships between the tissue structural constituents. Investigation reveals a novel complex cellular network in highly mineralized tendon aspects, where ∼100 nm diameter canaliculi emanating from cell (tenocyte) lacunae surround extracellular collagen fibril bundles. Canaliculi are linked to smaller channels of ∼40 nm diameter, occupying spaces between fibrils. Close to the tendon mineralization front, calcium-rich globules appear between the fibrils and, with time, mineral propagates along and within collagen. These close associations between tenocytes, canaliculi, small channels, collagen and mineral suggest a new concept for the mineralization process, where ions and/or mineral precursors may be transported through spaces between fibrils before they crystallize along the surface of and within the fibrils.

**Significance Statement:** The basic mechanism by which vertebrate collagenous tissues are mineralized is still not fully elucidated, despite the importance of this process for skeletal formation and regeneration. Through three-dimensional imaging of the cellular network together with the extracellular matrix and mineral deposits, the present work investigates normally mineralizing avian leg tendon as a model system for vertebrates in general. The data support a mechanism where mineral ions and possible mineral precursors are initially present in interfibrillar collagen spaces and are subsequently translocated to neighboring collagen fibrils. Mineral particles then nucleate in association with collagen to form the well known collagen-mineral composite material of the skeleton.

## Introduction

Bone and other collagen-based mineralized tissues such as dentin or mineralized tendon are materials remarkable for their hierarchical structure based on a unique combination of protein, mineral and water that provide relatively high fracture resistance (1, 2). Such mechanical competence implies not only a well-balanced ratio between its content of mineral and collagen in extracellular matrices (ECMs) but also appropriate spatial relations between all its components (3–5). Development of mineralized collagenous tissue is orchestrated in part by tissue-forming cells: osteoblasts in bone, odontoblasts in dentin, chondrocytes in cartilage and tenocytes in tendon synthesize and secrete collagen as a principal product together with non-collagenous proteins and other molecules that comprise ECMs where mineral deposition predominantly occurs (6–9). Osteocytes, and presumably certain of the other cell types above, generate and extend a series of processes that interconnect cells, conduct fluid, and provide essential means for cell-cell and cell-matrix crosstalk and communication (10, 11).

In bone, some osteoblasts differentiate to osteocytes and form a complex three-dimensional (3D) network where cells are connected through canaliculi across the mineralized tissue, which is known to be related to the microarchitecture of collagenous tissue, most notably the local fiber orientation (10, 12). Osteocyte networks have a variety of functions principally associated with mineral homeostasis in the body (13, 14). During bone remodeling, osteocyte networks are visible in newly deposited osteoid, (15) an observation which has led to the hypothesis that they may contribute to transport of mineralization precursors to the mineralization front (16, 17).

While the structure of mineralized collagen fibrils has long been studied by transmission electron microscopy and more recently by electron tomography (18–20), surprisingly little is known about the 3D arrangement of collagen fibrils and mineral with respect to the cell network within a mineralizing bone matrix. Such information is difficult to obtain because it requires nanometer resolution and a large field of view in the range of tens of microns, in order to visualize several cells with their processes (21). In this study, focused ion beam-scanning electron microscopy (FIB-SEM) (22) was used to examine simultaneously cells, ECM, and mineral deposits in turkey leg tendon (TLT) as a model system for collagen mineralization. Normally mineralizing tendons are representative of other vertebrate mineralizing tissues and are structurally more advantageous for study and analysis (23, 24). TLT is comprised of collagen fibrils elaborated in essentially parallel arrays rather than in more complex overlapping or twisted assemblages found in bone and other vertebrate systems (25). Furthermore, the sequence of events of mineral deposition may be followed and described spatially and temporally along the tendon length because the mineralization begins in the distal aspect of the tissue and continues uninterrupted proximally from that origin. These tissue characteristics have provided insight into certain aspects of the basic mechanism(s) of vertebrate mineralization and avian tendon adaptation in material properties (23, 24, 26).

In fully mineralized tendon, a previously unknown extensive 3D network of tenocytes-canaliculi resembling the network of osteocytes-canaliculi in bone was found. This network branches into finer channels that appear to occupy spaces between collagen fibrils. At the tendon mineralization front, strings of calcium-rich globules were observed in such channels between fibrils which subsequently propagate into neighboring fibrils. Collectively, these data begin to define possible temporal and spatial relationships between tendon cells, their elaborated ECM, and both inter- and intra-fibrillar collagen mineralization as well as further understanding of critical events mediating mineralization in vertebrate tissues.

## Results and Discussion

### Tenocyte networks in the mineralized zone of turkey leg tendon

We first investigated the fully mineralized zone of the tibialis cranialis tendon from the 26-week-old male domestic turkey, *Meleagris gallopavo*, across multiple length scales. Micro-computed tomography (micro-CT) of the distal tendon (D) showed a clear contrast between its radiopaque mineralized region (M) and dark grey unmineralized zone (UM) (Fig. 1a). Large canals ∼100 µm in diameter were clearly visible in the mineralized zone (Fig. 1b, pink; Fig. S1a-d), oriented mainly along the longitudinal direction of the tendon and interconnected through oblique perforating channels. Transverse sections of the fully mineralized zone (Fig. 1c) revealed two types of tissues: circumferential (CIR) tissue with lower porosity surrounding the canals and interstitial (INT) tissue with higher porosity occupying the spaces between CIR zones. In agreement with a previous report (27), the volume fraction of CIR tissue qualitatively decreased from the more highly mineralized (mature) tendon region to the region closer to the mineralization front (Movie S1). Interestingly, the CIR tissue together with the canals resembles the morphology of Haversian systems in bone (28) and the central canals may contain blood vessels to provide ions, nutrients and other substituents for cell viability, which may play a critical role in the mineralization process.

**Fig. 1:**
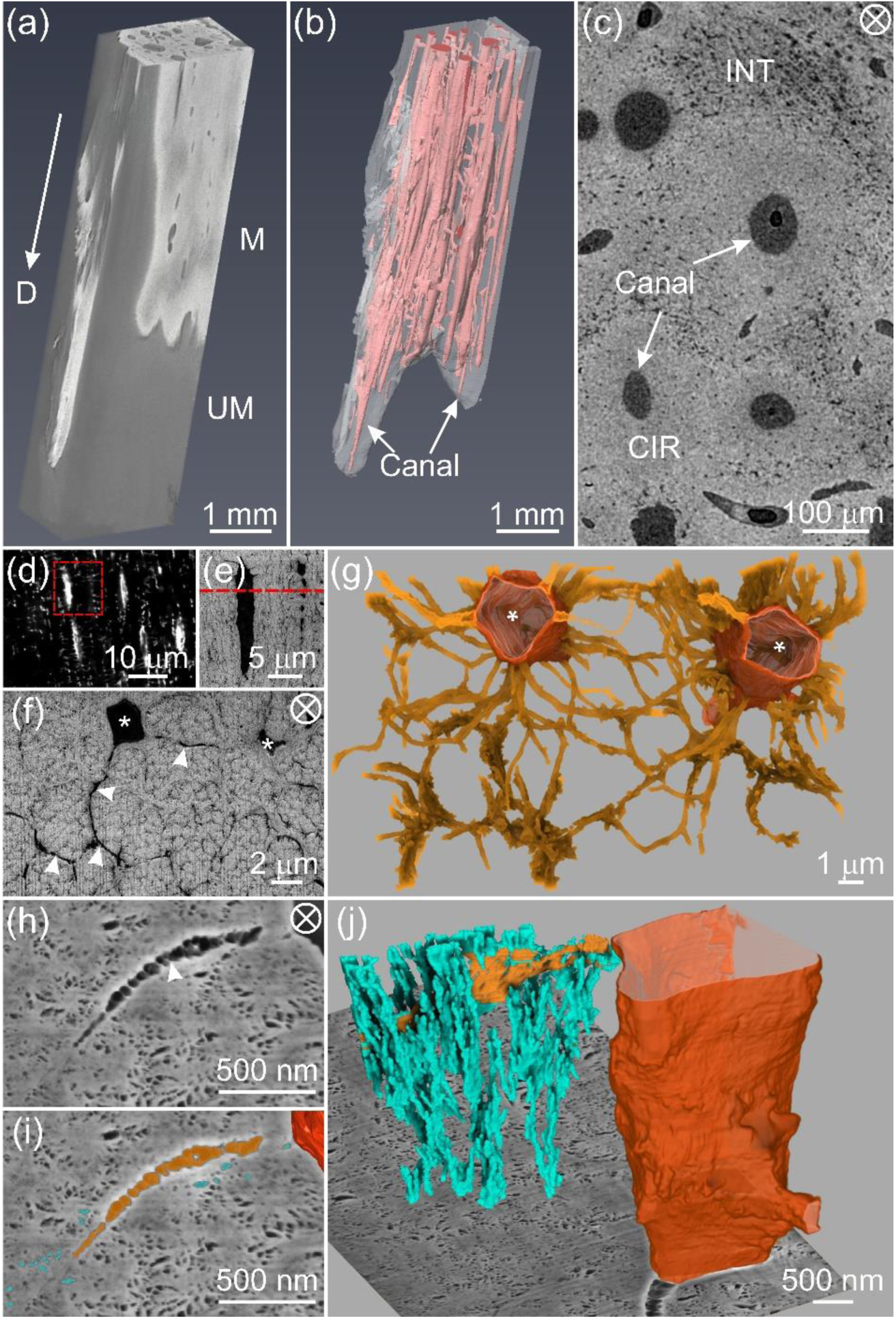
Tenocyte network in a highly mineralized zone of TLT from a tendon prepared by chemical fixation and stained with osmium tetroxide alone. **a**, 3D reconstruction of micro-CT data from the more heavily mineralized distal end of the leg tendon. This region contains both mineralized (radiopaque, M) and unmineralized (darker grey, UM) zones. The arrow points toward the distal end (D) of the tendon. **b**, The mineralized zone in **a** contains multiple unmineralized canals (pink) principally oriented in the tendon longitudinal direction. **c**, High resolution micro-CT image of a representative transverse section from a fully mineralized zone of the tendon reveals circumferential regions (CIR) with qualitatively lower porosity surrounding the canals and interstitial tissue (INT) with higher porosity between CIRs. **d**, Confocal image of a longitudinal section in the mineralized zone of a Rhodamine-stained tendon showing tenocyte lacunae and canaliculi (white streaks and small white dots and lines, respectively). An area framed in red is enlarged and shown by BSE imaging in **e** where lacunae and canaliculi are more apparent. **f**, BSE image of a transverse section (marked by the red dashed line in **e**) showing two tenocyte lacunae (*) and radiating canaliculi (arrowheads) surrounding collagen fibril bundles. **g**, FIB-SEM 3D reconstruction of a volume of **f** illustrating the two tenocyte lacunae (tangerine) and their associated canaliculi (gold). **h**, High resolution SE image of a tendon section showing a portion of a cell lacuna, canaliculus (arrowhead) and fine pores, highlighted in dark tangerine, gold, and turquoise, respectively in **i. j**, 3D rendering corresponding to **h, i** of the tenocyte lacuna-canalicular network intersecting with a FIB-SEM background image plane and demonstrating the numerous secondary channels (turquoise) branching from a canaliculus (gold) and disposed primarily in the longitudinal tendon direction. The tenocyte lacuna is defined by a contoured surface (dark tangerine). The circled cross in **c, f**, and **h** denotes the view along the longitudinal direction of the tendon.

Reconstruction of micro-CT images (Fig. S1e-h) revealed elongated/prolate cell lacunae in the CIR regions principally oriented along the tendon long axis. At higher resolution, laser scanning confocal microscopy (LSCM) imaging (Fig. 1d, Fig. S2 and Movie S2) demonstrated a cellular network that is similar to the lacuno-canalicular network in bone(10): the spindle-shaped cell lacunae were oriented along the longitudinal direction of the tendon and the canaliculi were radiating from the lacunae, mainly perpendicular to the lacunae long axes. The same area was further investigated by FIB-SEM imaging with a voxel size of (10.6 × 10.6 × 21) nm^3^ (Fig. 1e-j and Movie S3), which showed the canaliculi with an average diameter of ∼100 nm, similar to that found in mouse femoral bone (29), surrounding and occasionally traversing collagen fibril bundles (Fig. 1g).

High resolution FIB-SEM imaging with a voxel size of (6.0 × 6.0 × 8.7) nm^3^ (Fig. 1h-j and Movie S4) revealed a significant number of finer pores with a diameter of ∼40 nm connected to the canaliculi. 3D reconstruction of these pores showed that they formed channels (Fig. 1i and j) predominantly along the longitudinal direction of collagen fibril bundles. Whether these pores are filled with fluid or may contain organic material such as glycosaminoglycans and/or cell processes is difficult to determine with current FIB-SEM analysis. Nonetheless, these smaller interconnected pores, termed secondary channels, might facilitate the mineralization of collagen fibrils that are far from the tenocytes of the tissue through possible mechanisms of transport or diffusion of ions or mineral precursors, for example.

### Tenocytes, collagen fibrils and mineral deposits at the mineralization front

To gain insight into the process of vertebrate mineralization, thin longitudinal sections (∼150 µm) of leg tendons from freshly sacrificed 23-week-old turkeys were investigated by FIB-SEM. The tissues contained the transitional region between unmineralized and mineralized portions of the tendon and were prepared by high pressure freezing (HPF) followed by automated freeze substitution (AFS) with uranyl acetate staining. Tenocytes, ECM components, particularly collagen fibril bundles, and mineral deposits could be clearly identified in SEM images (Fig. 2a). From longitudinal views with respect to the tendon long axis (Fig. 2b), for example, four tenocytes were seen aligned along collagen fibril bundles. Such alignment could be more clearly visualized by 3D rendering where tenocytes were in close contact with mineralized collagen fibril bundles (Fig. 2c and Movie S5). The bundles were circumferentially enclosed by a thin sheath, known as the endotendon (9), and themselves contained individual mineralized collagen fibrils (Fig. S3). Such sheaths enclosing collagen bundles were also observed in unmineralized tendon regions, where cells and their extensions were situated in the spaces between several collagen fibril bundles (Fig. S4). A recent mouse tendon study by Kalson et al. (30) demonstrated similar results where cells had elongated and interconnected cell processes, which delimited collagen fibril bundles. These data suggest that cell processes may overlap and/or may be embedded in the sheath surrounding collagen fibril bundles. While collagen becomes mineralized at later stages of tendon development, the canaliculi associated with sheath regions remain unmineralized and may serve to accommodate cell processes when the tissue becomes more completely mineralized (Fig. S4).

**Fig. 2:**
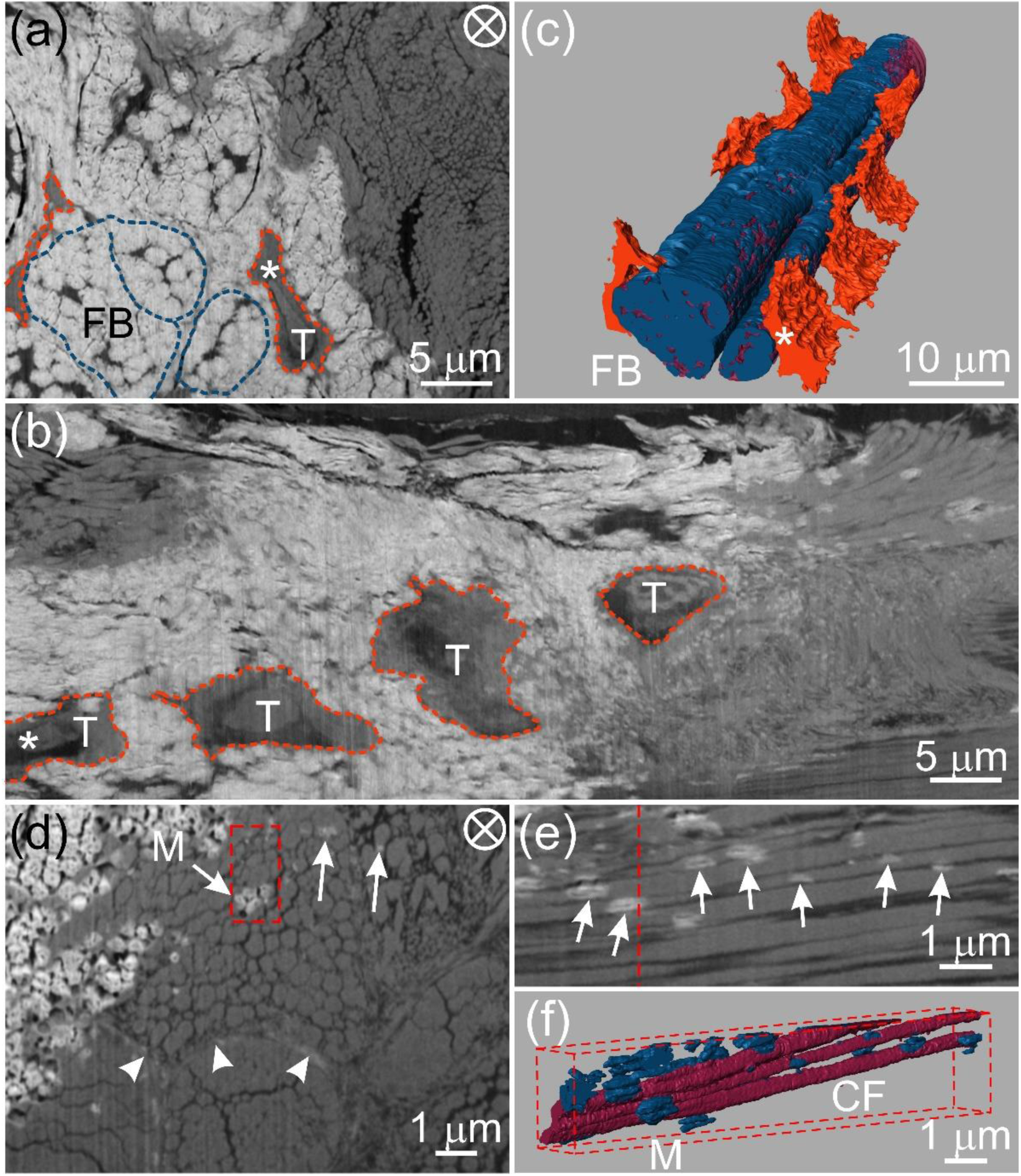
FIB-SEM images and corresponding 3D reconstruction of tenocytes, collagen fibrils and mineral deposits at an interface between mineralized and unmineralized avian tendon zones. Images are from the same specimen, prepared by cryomicrotomy and HPF. AFS followed these procedures and the sample was stained with uranyl acetate alone. **a**, A BSE image of a transverse section of a typical tendon volume showing mineralized (white) and unmineralized (grey) zones. Two collagen fibril bundles (FB) and a single tenocyte (T) may be identified in the mineralized zone and are denoted by dashed blue and tangerine lines, respectively. There are thin, dark structures of unknown nature that appear to separate individual mineralized units within bundles. Such structures may possibly be related to canaliculi surrounding fibril bundles. **b**, Longitudinal view obtained by digital processing the full stack of FIB-SEM slices through the same tendon volume represented by the single transverse BSE image in **a. c**, 3D reconstruction of the FIB-SEM data of the volume of **a** and **b** showing tenocytes (tangerine) adjacent to and aligned along the longitudinal direction of mineralized collagen fibril bundles. Bundles may be distinguished by their component collagen fibrils (wine) and mineral (blue). **d**, High resolution BSE image showing mineral (arrows, M) within collagen fibril bundles and sheath regions (arrowheads). **e**, Longitudinal view obtained by digital processing the full stack of FIB-SEM slices through the same tendon volume represented by the transverse BSE image in the red framed area of **d**. The red dashed line in **e** represents the location of the framed area of **d**. Numerous mineral deposits (arrows) are associated with collagen fibrils and appear predominantly as prolate ellipsoids in shape and elongated in the tendon longitudinal direction. **f**, 3D reconstruction of the volume marked by red framed area in **d** showing several collagen fibrils (CF, wine) and their associated discrete mineral deposits (M, blue). As in **e**, the longitudinal view reveals numerous mineral deposits interrelated with several collagen fibrils, many of which are separated by dark, narrow spaces. The circled cross in **a, d** denotes the view along the longitudinal direction of the tendon.

High resolution FIB-SEM data with a voxel size of (12.0 × 12.0 × 24.0) nm^3^ revealed isolated mineral deposits far from tenocytes and within both collagen fibril bundles (arrows, Fig. 2d) and sheath regions (arrowheads, Fig. 2d). Mineral deposits within collagen fibril bundles framed in Fig. 2d were digitally processed in a longitudinal view (Fig. 2e) and rendered three-dimensionally in a slightly oblique view (Fig. 2f). Such longitudinal perspectives clearly demonstrate that the mineral deposits (arrows, Fig. 2e; blue, Fig. 2f) have a prolate ellipsoidal shape elongated along the tendon longitudinal direction. This shape suggests that the minerals grow faster along the longitudinal rather than the radial direction of collagen fibrils. This observation is consistent with reports that mineral crystals grow preferentially along their crystallographic c-axes and in the direction of the collagen long axis (31). Fig. 2e also shows spaces averaging ∼40 nm in width between adjacent collagen fibrils viewed in longitudinal profile. Such spaces may correspond to the secondary channels identified and described in Fig. 1f.

To investigate the spatial relationships between collagen fibrils and mineral deposits and to understand the progression of mineralization events, tendons from 23-week-old turkeys were cryo-fixed utilizing HPF, stained with osmium tetroxide and uranyl acetate and examined ultrastructurally with FIB-SEM with a voxel size of (6.0 × 6.0 × 8.7) nm^3^. Both transverse (Fig. 3a-b) and longitudinal (Fig. 4d-f) sections of tendons showed relatively thinner collagen fibrils (∼30-50 nm in diameter) joining or fusing with thicker fibrils (∼200-300 nm in diameter). Thinner fibrils represent more recently synthesized and secreted collagen compared to thicker fibrils. Both thinner and thicker fibrils maintain the characteristic collagen periodicity (Fig. 4f). In many instances, such thinner fibrils, as well thicker fibrils, were found in the vicinity of the earliest detectable mineral deposits in tendon regions (Fig. 3a and 4d, f). Deposits may come into close contact with both thin and thick fibrils (Fig. 4d, f) and their disposition between fibrils of any size constitutes interfibrillar mineral. Deposits located within fibrils define intrafibrillar mineral.

**Fig. 3:**
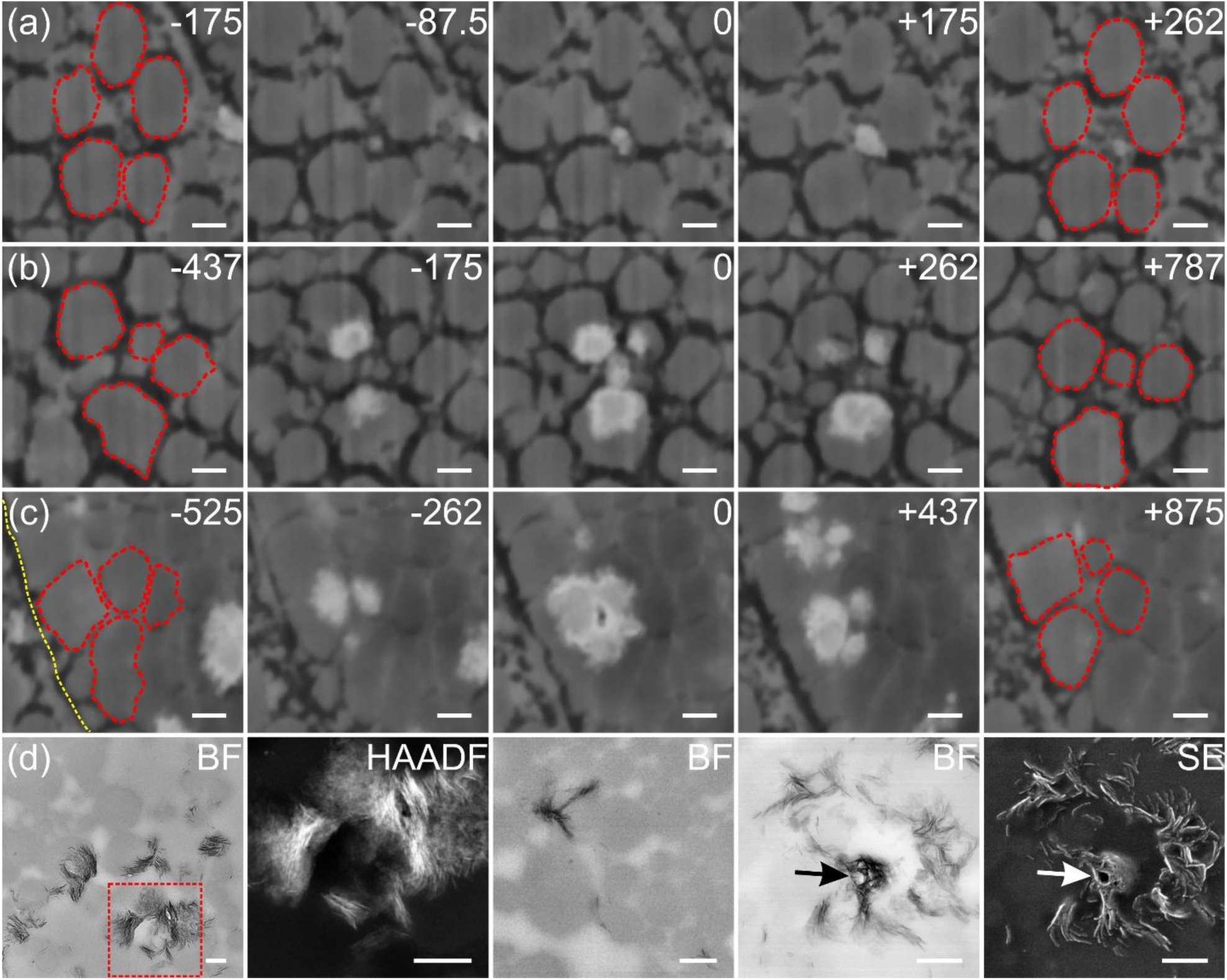
Series of FIB-SEM (panels a-c) and TEM images (panel d) showing spatial relationships between collagen fibrils (red outlines) and mineral deposits (white) in a distal region of an avian tendon. The FIB-SEM panels show different tissue regions in transverse profile in a single tendon specimen, prepared by HPF followed by AFS and staining with osmium tetroxide-uranyl acetate. Each FIB-SEM panel of five images represents mineralization occurring at different degrees at the specific locations imaged. Numbers at the top right corners of panels **a**-**c** represent the distance (nm) of each image to the reference image (marked by 0) along the longitudinal direction of the collagen fibrils (the tendon long axis) in the specimen volume investigated: **a**, Early stage mineral formation beginning and developing among five unmineralized collagen fibrils. **b**, Several mineral deposits appearing principally as intrafibrillar collagen mineralization. The deposits of mineral occupy only a portion of the fibrils and are located at their peripheries. **c**, A region of densely packed collagen fibrils illustrating a degree of intrafibrillar mineralization that reveals a small dark area, possibly a narrow channel, in the central interfibrillar space. The yellow dotted line (panel 1 from the left of Fig. 3c) represents the boundary of the collagen fibril bundle containing the fibrils of interest. **d**, Bright field (BF, far left panel) and high angle annular dark field (HAADF) images of a tendon sample prepared by chemical fixation. The images show mineral deposits principally surrounding collagen fibrils as viewed in transverse profiles. The HAADF image was obtained from the red-framed region of BF (far left panel). The middle BF image demonstrates intrafibrillar mineral deposits (black) spanning several collagen fibrils (grey). BF and SE images (panels 4-5, respectively, from the left) show a possible channel (arrows) encircled by mineral in interfibrillar space. Scale bar: 200 nm for all panels.

**Fig. 4:**
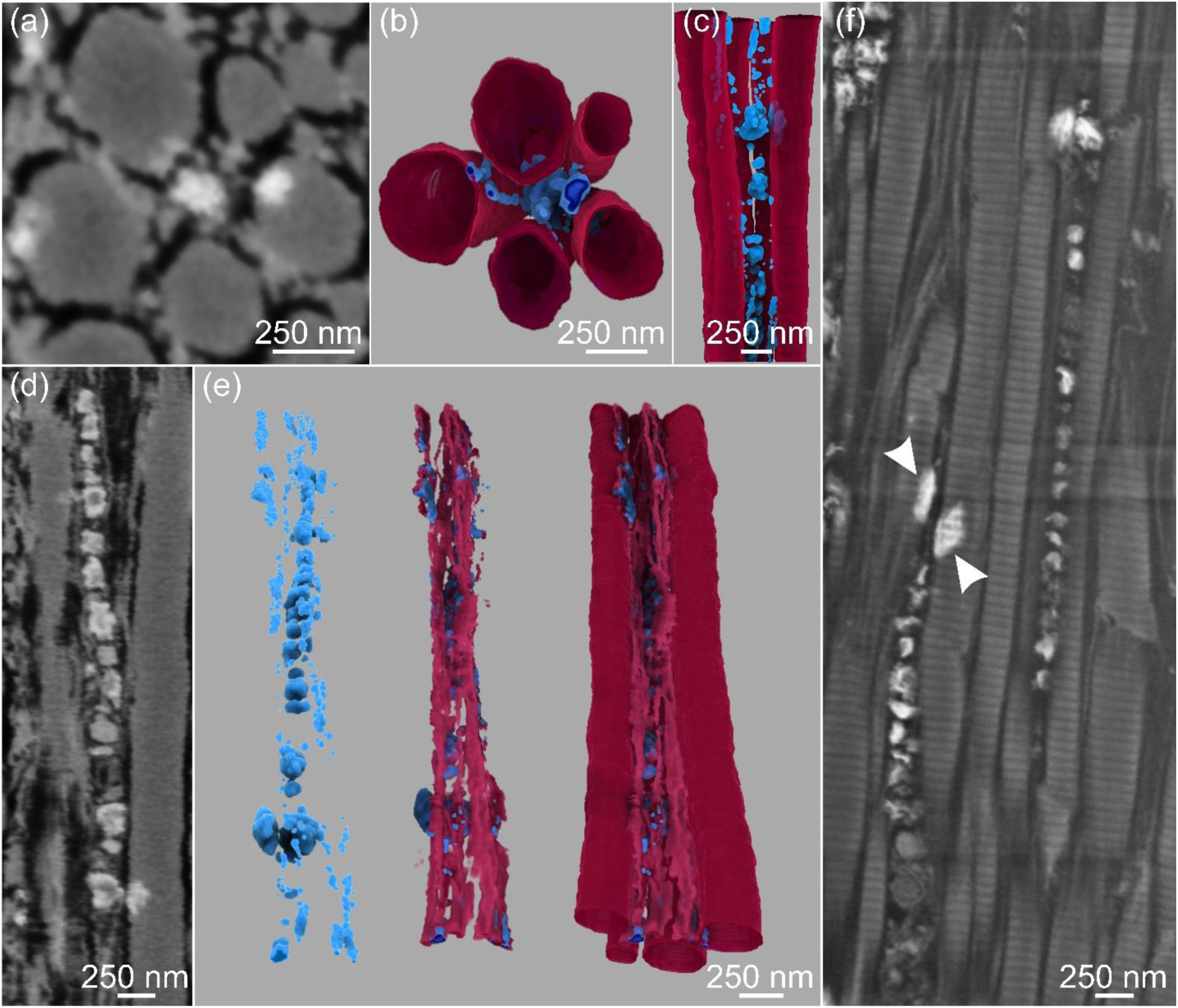
FIB-SEM images (a, d and f) and 3D reconstructions (b, c and e) of early stages of mineralization showing structural and spatial relationships between mineral deposits, possible secondary channels, and collagen fibrils in avian tendon. The tendon specimen was prepared by HPF followed by AFS and staining with osmium tetroxide-uranyl acetate. **a**-**c, d**-**e**, and **f** represent three different regions of the same tendon, distal (**a-c** and **d-e**) and proximal (**f**). **a**, SE image of a transverse section of five collagen fibrils, the intervening spaces between them (black) which may be secondary channels, and several small mineral deposits (white). **b**, transverse, top-down, and **c**, longitudinal view of the 3D reconstruction of the volume represented by **a**, showing the mineral deposits (blue) both within and outside the collagen fibrils (wine). **d**, SE image of the longitudinal section illustrates linearly arranged mineral deposits (white) of different sizes predominantly between thin and thick collagen fibrils in possible secondary channels (black). **e**, 3D reconstructions of **d** showing mineral deposits only (blue; left image), mineral associated with thin collagen fibrils (light wine; middle image), and mineral associated with both thin and thick fibrils (wine; right image) with mineral deposits aligned along the longitudinal collagen axes. **f**, High resolution SE image of a region of tendon ECM different from that in **d** but showing similar collagen mineralization characteristics; periodic banding of both thin and thick collagen fibrils is clear, thin and thick fibrils fuse together, and mineral (white) is found in linear disposition between fibrils as well as on the surfaces and within fibrils (arrowheads). The mineral may be associated with matrix vesicles as noted previously although such a relationship has not been established definitively. Spaces (black) between fibrils may denote putative secondary channels in the tendon matrix.

Along the tendon long axis, successive FIB slices represent progressive changes in tissue location, time and degree of tendon mineralization, and corresponding SEM images showed that mineral deposits grew both along and within collagen fibrils (inter- and intra-fibrillar collagen mineralization, respectively) (Fig. 3b and Movie S6). In some aspects of mineralization, possible channels ∼40 nm in diameter were observed in the central space within a group of mineralized fibrils (Fig. 3c). Such spaces may correspond to the dark, narrow separations found in longitudinal sections of tendon (Fig. 2e). TEM analysis of similar mineralizing tendon regions (Fig. 3d) also indicated the presence of plate-like mineral crystals within and outside collagen fibrils. Further, these mineral crystals were found to encircle channel-like structures (arrows) observed in bright field and secondary electron images (panels 4-5 from the left of Fig. 3d), results consistent with FIB-SEM findings (Fig. 2e, 3c).

Whether collagen fibril mineralization occurs in an inter- or intra-fibrillar manner is critical to understanding mineralization mechanisms in vertebrates in general. Recently, by using FIB-SEM and TEM imaging, Reznikov et al. (20) suggested that mineral deposits in bone form an interlinked network through cross-fibrillar mineralization, that is, by both inter- and intra-fibrillar mineralization. This concept was implicit and proposed in earlier work with avian tendon examined by TEM (23, 32, 33). Here, segmentation and 3D reconstruction of tendon collagen fibrils and mineral deposits provided new insight into collagen-mineral interactions regarding inter- and intra-fibrillar mineralization of this tissue (Fig. 4). Firstly, numerous mineral deposits of variable sizes clearly appeared both between and within collagen fibrils. Fibrils, themselves, were of different and variable diameters, as noted above. The individual mineral deposits (for example, Fig. 4a, d, f), rich in calcium and phosphate as detected by energy dispersive X-ray spectroscopy (EDS) (Fig. S5), appeared to be located between collagen fibrils, whether thin or thick (interfibrillar mineral), as well as within thick fibrils (intrafibrillar mineral) (Fig. 4f). Deposits furthermore were frequently found in linear arrangements in interfibrillar spaces (Fig. 4c-f) and, as such, were strikingly similar to those reported in previous TEM studies (23, 33) of normally mineralizing avian tendon. The deposits were suggested to be associated with extracellular vesicles disposed in spaces between collagen fibrils, (23, 33) and the vesicles appeared to maintain a close spatial relation with the extensive network of tenocyte cell processes in the tissue (33).

In the present investigation, extracellular vesicles could not be directly visualized since their investing membranes were not entirely clear. Previous studies report that vesicles undergo changes resulting, for example, in the degradation of their membranes and the subsequent loss or release of their mineral contents to the ECM milieu (34, 35). Also because their membranes were uncertain, cell processes have not been directly identified in the current work, but it may be assumed that they occupy the spaces defined by canaliculi and secondary channels, as mentioned above. Thus, the tendon establishes and maintains an ultrastructural organization of collagen and spatially interrelated cell processes, ECM vesicles and mineral deposits. In this regard, the processes, vesicles and deposits provide a pathway by which mineral ions and/or mineral precursors may be actively transported, become accessible to collagen, and lead to collagen mineralization. Further, the image data of the present work (Fig. 3, for example) support the concept that interfibrillar mineral deposition precedes intrafibrillar mineralization, a result that is consistent with previous reports (32, 36).

The work here has also shown mineral deposition associated with the surfaces and intrafibrillar domains of collagen (Fig. 4f). Such mineral formation within collagen commonly occurred at sites near every or nearly every putative mineralizing vesicle noted above in the interfibrillar collagen spaces (Fig. 4f and Movie S7). On reconstruction of mineral deposits in a representative volume of the ECM, the interfibrillar mineralized sites were isolated from each other and much more numerous than could be revealed in individual 2D SEM slices (Movie S7). It also appeared as if individual interfibrillar mineral deposits were separated by a similar distance as they formed in linear arrangements (Movie S7). No direct interconnection can be clearly detected between inter- and intra-fibrillar mineralization sites, so an immediate structural relationship, if any, between them cannot be determined. In interfibrillar spaces, it may be reasonable to assume that local changes occur in the concentrations of mineral ions or possible mineral precursors resulting from their loss or release from vesicles as noted above. Such fluctuations in mineral ion or precursor concentrations then alter the physicochemical metastability of fluid ion concentration surrounding collagen and mediate formation and propagation of mineral over collagen surfaces and/or within collagen intrafibrillar domains. In addition, the establishment of simultaneous electrostatic neutrality and osmotic equilibrium between inter- and intra-fibrillar spaces of collagen, as described by the Gibbs-Donnan model (37), has been proposed recently to explain the migration of mineral ions or mineral precursors into intrafibrillar collagen spaces (38). Thus, the normally mineralizing tendon model illustrates both inter- and intra-fibrillar mineral deposition and yields results consistent with those of bone and tendon as described previously (20, 23, 32, 39).

The presence of a canalicular network throughout the ECM of avian tendon is compatible with the idea that mineral ions may be passively transported in its fluid-filled cavities. In this regard, the possibility exists that such mineral ions move about the interfibrillar collagen spaces and may mediate mineral deposition and its progression. This mechanism of passive transport may be independent or complementary to that provided by the putative active transport of ions and/or mineral precursors through vesicles described previously. Whatever the situation, the space between collagen fibrils appears to be critical for ion transport leading to mineral deposition of those fibrils comprising tendon structure.

Nucleation of mineral in association with collagen surfaces or within collagen fibrils has been conceptualized through different mechanisms. One such proposal involves calcium ion transport to collagen by binding to a negatively charged polyelectrolyte, such as phosphorylated osteopontin (40). Alternatively, ions may diffuse or be actively or passively transported as described above. In either case, the mineral ions are thought to associate and bind to specific sites defined stereochemically by charged amino acid residues comprising collagen α-chains located at the surfaces or within collagen fibrils (36, 41). Other potential sites of mineral ion binding at the collagen surfaces may be provided by charged, non-collagenous proteins such as fetuin, bone sialoprotein, osteocalcin and osteopontin, which themselves are surface-bound to collagen (36, 42–44). Additionally, osteocalcin is known to reside within collagen and in its so-called hole zones where it might bind calcium to facilitate nucleation (42).

Based on the FIB-SEM observations from tendon mineralization fronts and fully mineralized zones, the spatial interrelations between tenocytes, their ECM and mineral deposits are proposed and schematically shown in Fig. 5. Tenocytes are interconnected predominantly through canaliculi, which surround each collagen fibril bundle circumferentially and occasionally traverse through them (Fig. S4). Numerous finer channels branch predominantly in orthogonal directions from the canaliculi and pervade collagen fibril bundles approximately parallel to the fibril bundle longitudinal axes. The extent of such secondary channels intersecting with collagen fibril bundles has not yet been fully documented here by FIB-SEM and 3D reconstruction but, from initial data as represented by Fig. 1j and earlier 2D TEM images, (23, 33) it is likely that secondary channels branch at multiple points and provide fluid and mineral ion access through cell processes and matrix vesicles to not only fibril bundle surfaces but also individual collagen fibrils within bundles. The nature of this suggested multiple branching and the direction of secondary channels with respect to bundles and fibrils can only be speculative at this point. Fibril bundles consist of numerous collagen fibrils that are separated by the secondary channels, and mineral deposition occurs in the channel spaces as well as on the surfaces and within fibrils. Potentially, these interfibrillar secondary channels could allow the infiltration of the collagen matrix by many different ions, for example, calcium ions electrostatically bound to phosphorylated proteins that serve at the same time as calcium transporters and crystallization inhibitors, as proposed from mineralization studies in vitro, while mineral nucleation occurs on the surfaces of and within collagen fibrils (45–47).

**Fig. 5:**
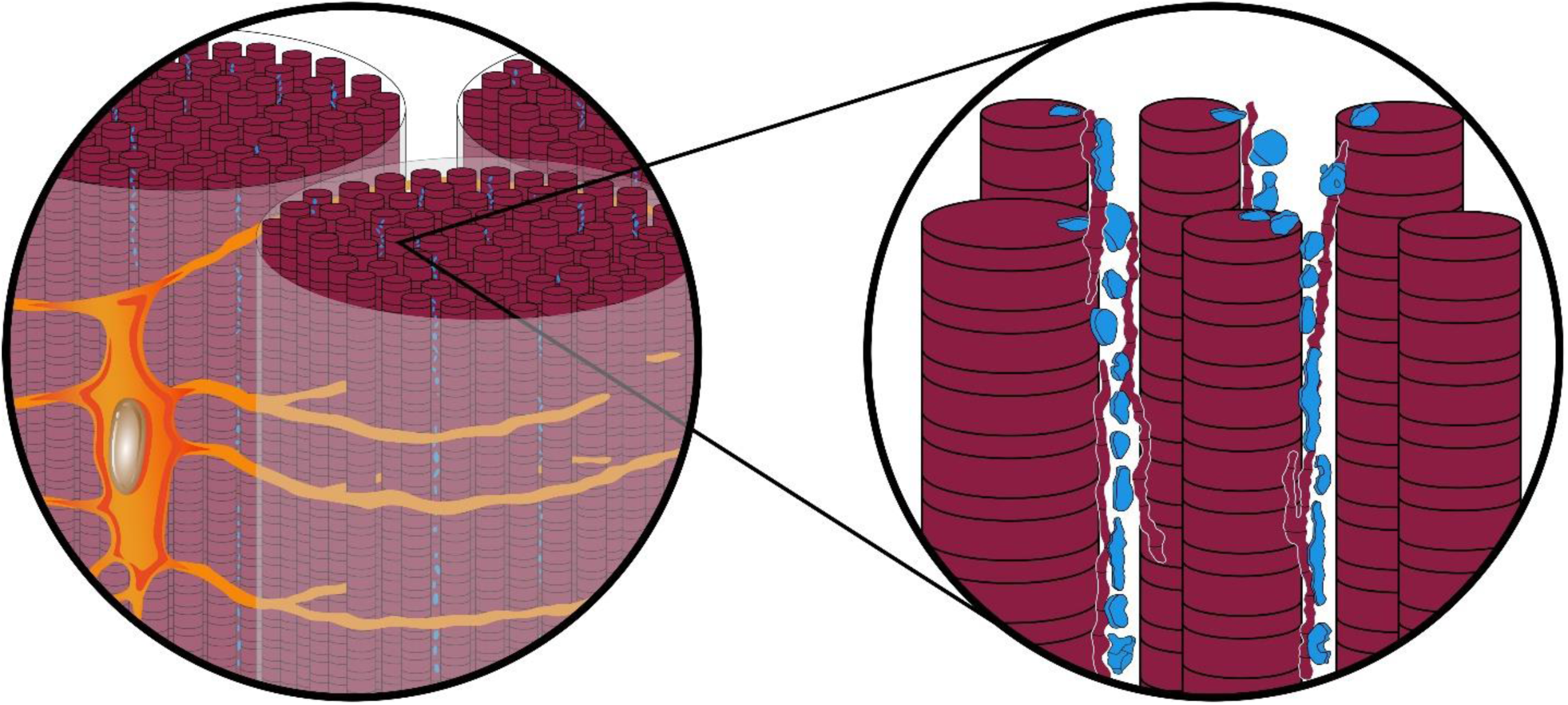
Schematic illustration showing proposed ultrastructural interrelationships between tenocytes, their secreted ECM and mineral in avian leg tendon. Sections of tissue viewed along the long axis of collagen fibril bundles (left panel) show a single tenocyte (tangerine) with its nucleus (grey-brown) and a number of cell processes (amber). Processes enter canaliculi (gold) that envelope fibril bundles about their surfaces and may branch into the bundle sheath (pale white). On enlargement (right panel), numerous collagen fibrils (wine) comprising bundles consist of both thinner and thicker structures separated by secondary channels (white spaces) which originate in orthogonal directions from canaliculi and along collagen long axes. Mineral deposits (blue) appear outside and along the surfaces (interfibrillar locations) as well as inside (intrafibrillar locations) collagen fibrils. Banding about collagen represents its characteristic periodicity observed by electron microscopy.

### Potential concerns, limitations and novelty of this study

While the data and concepts revealed and discussed above generate novel information regarding mineralization of vertebrate tissues, there are several areas of the study that remain uncertain in nature. A principal concern in general is the possible creation of microscopic artefact potentially the result of dehydration of tendon tissue during specimen processing. In this regard, there may be shrinkage of tissue that leads to spaces between structural components such as thin and thick collagen fibrils and the misleading creation of canaliculi and secondary channels in tendon, for example. Additionally, dehydration may effectively concentrate mineral ions to produce the small, vesicle-like structures reported between, on the surface of, and within collagen. On the other hand, HPF and AFS mitigate shrinkage and dehydration and optimize tissue preparation for microscopic analyses and, furthermore, collagen of various diameters and mineralizing matrix vesicles have been repeatedly documented in numerous previous studies in which tissues have been prepared by a variety of techniques, including conventional, anhydrous as well as cryo-fixed methods. The presence of such tissue components observed regardless of preparative techniques would tend to diminish concerns that they are artefactual. Additionally, the present study does not address the possible presence of cell processes that are suggested to reside within canaliculi and secondary channels of the tendon. Detection of such processes will be the subject of further investigation in mineralizing avian tendon. The means by which mineral is propagated from inter- to intra-fibrillar tissue spaces remain conjectural and elusive. This subject, too, will be investigated more completely in other work. Finally, there is some caution to this study in terms of its sample pool; here, the number of animals (three) and the type of tendon (tibialis cranialis from among many mineralizing leg tendons) examined in detail were limited and FIB-SEM data were relevant to only a selected few regions of interest. Nonetheless, structural information obtained regarding collagen, mineralizing matrix vesicles, and the presence of mineral deposits associated with these tissue constituents was again largely consistent with that of previous reports. Considering the highly consistent nature of earlier and present data, the current work has yielded the extremely novel observations of canaliculi and secondary channels and their apparent interconnections for the first time in 3D. This study has led further to new insight into possible mechanisms by which mineral ions in the supersaturated fluid bathing the tissue as well as potential mineral precursors may be directed and transported throughout the tendon ECM.

## Summary and Outlook

In summary, FIB-SEM tomographic imaging has revealed novel 3D ultrastructural interrelations between tenocytes, tenocyte networks, ECM, and mineral deposits in normally mineralizing avian leg tendon. A complex and extensive cellular network was found consisting of canaliculi and much finer secondary channels that enveloped collagen fibril bundles and penetrated their sheaths. Further, secondary channels separated collagen fibrils within bundles and putatively provided the space for cell processes and the fluid environment passageway for mineral ion and/or mineral precursor transport. Possible matrix vesicles, commonly in linear arrangements, may originate from cell processes and their mineralization outside collagen fibrils appeared to serve as a source of interfibrillar mineralization of the matrices. At subsequent stages of mineral formation, mineral deposits were observed on and along surfaces of collagen fibrils (interfibrillar mineralization) as well as within fibrils (intrafibrillar mineralization). Overall, this report advances knowledge and understanding of cellular and ECM architecture not previously appreciated from 2D image analyses and yields new insight into the mechanism(s) of mineralization in bone, dentin, cementum, and other vertebrate tissues. Additional studies of normally mineralizing avian tendon and its counterpart vertebrate tissues to elaborate secondary channel ultrastructure, the nature of both passive and active transport, and the precise means of mineral propagation outside and within collagen will address aspects of mineral deposition that still remain uncertain.

## Materials and Methods

### Specimen preparation

Three 23-week-old and two 26-week-old male domestic turkeys, species *Meleagris gallopavo*, were obtained from local farms (Ullrichs Putenhof, Baden-Wurttemberg, and Hartmanns Putenhof, Bavaria, Germany) and freshly sacrificed. The tibialis cranialis tendons, containing the transitional region between noncalcified and calcified portions of the tissue, were carefully dissected from turkey legs, and the muscles attached to the tendons were removed using a scalpel. Dissection was conducted on an ice bath of phosphate-buffered saline solution (PBS; Sigma-Aldrich, St. Louis, MO, USA). The tendons were then either sectioned with a cryomicrotome (HM 560 CryoStar Cryostat, Thermo Scientific, Waltham, MA, USA) or cut with a scalpel to obtain thin sections 150-200 μm thick, followed by HPF. More specifically, samples were sandwiched between two carriers (200 μm brass planchet, Type B freezer hats, Ted Pella, Inc., Redding, CA, USA), filled with 1-hexadecane (Sigma-Aldrich) as cryoprotectant and then cryofixed in a model Leica EM HPM100 high pressure freezing machine (Leica Microsystems, Wetzlar, Germany).

Freeze substitution was carried out in a Leica EM AFS2 automated freeze-substitution unit (Leica Microsystems). The samples, still attached to the frozen carriers, were transferred in liquid nitrogen to 1 mL pre-cooled freeze-substitution media (at −90 °C) in cryo-vials. The following freeze-substitution protocols were used to enhance the imaging contrast of unmineralized tissue for focused ion beam-scanning electron microscopy (FIB-SEM) and transmission electron microscopy (TEM):

#### Protocol 1

The freeze-substitution media consisted of 0.2% (w/v) uranyl acetate (SERVA Electrophoresis GmbH, Heidelberg, Germany) in absolute acetone (diluted from 20% uranyl acetate stock in methanol (Sigma-Aldrich)). The temperature of the AFS processing chamber was first maintained at −85 °C for 16 h, then raised to −50 °C at a rate of 7 °C/h, held at −50 °C for 7 h, raised to 0 °C at a rate of 5 °C/h and maintained at 0 °C for 72 h. When the temperature was −50 °C (third step as mentioned above), the sample carriers were removed in liquid nitrogen and the samples were stepwise infiltrated by Lowicryl HM20 (Polysciences, Inc., Warrington, PA, USA) with a progressively increasing ratio of HM20 to acetone (volume fraction of HM20: 0%, 30%, 50%, 70% and 100%; 1 h for each step). Finally, the samples were transferred to fresh 100% HM20. Polymerization under UV light was begun at the fourth step of infiltration and continued to the end of the procedure.

#### Protocol 2

A mixture of 0.1% osmium tetroxide (Electron Microscopy Sciences, Hatfield, PA, USA), 0.1% uranyl acetate (SERVA Electrophoresis), 0.5% glutaraldehyde (Electron Microscopy Sciences), 1.5% H_2_O and 100% acetone was used for freeze substitution. The AFS temperature progression began at −120 °C for 3 h, followed by warming to −85 °C at a rate of 17.5 °C/h, maintaining −85 °C for 103 h, warming to −20 °C at a rate of 7.3 °C/h, maintaining −20 °C for 12 h and finally warming to 4 °C at a rate of 2.6 °C/h. At this point, the samples were removed from the specimen carriers (at room temperature) and subsequently embedded in EPON resin (Polysciences, Inc.).

Conventional chemical fixation of freshly acquired samples was also used. After dissection of the turkey leg tendon, samples were chemically fixed in 4% paraformaldehyde (Electron Microscopy Sciences) and 2% glutaraldehyde (Electron Microscopy Sciences) in cacodylate buffer solution (Electron Microscopy Sciences) for 8 h, followed by staining with 2% osmium tetroxide (Electron Microscopy Sciences) in PBS solution for 2 h. After washing with PBS solution, the samples were dehydrated in a graded acetone series (30%, 50%, 70%, 90%, 100% and 100%; 1 h for each step) and then stained with 0.004% (wt/v) Rhodamine 6G (Thermo Fisher) in acetone and finally embedded in Spurr resin (Polysciences, Inc.).

All embedded samples were ground with a series of carbide grinding papers to expose longitudinal sections of turkey leg tendons and then polished with a diamond suspension to improve surface smoothness for subsequent serial FIB-SEM imaging.

### Micro-computed tomography

Spurr-embedded tendon sections containing both mineralized and unmineralized regions of the tissues were imaged using a Micro-CT imager (RX Solutions EasyTom160/150 tomographic unit, Chavanod, France) at a resolution of 4.5 µm/voxel for larger volume observations and 1.0 µm/voxel for more refined volume observations at 100 kV and 200 µA. X-Act CT software (RX Solutions) was used to reconstruct the projection images.

### Laser scanning confocal microscopy

A Leica TCS SP8 laser scanning confocal microscope (LSCM; Leica Microsystems) with a 40X/1.3 oil objective lens was used to identify tendon cells, organization and connections through canaliculi. The 526 nm line of a multi-line Argon ion laser was used for Rhodamine 6G excitation with emission at 555 nm. Selected sites of mineralized tissue were further characterized in 3D, during which the specimens were typically imaged from the surface to depths of 30 µm with a step size of 300 nm for each image. Each sequence of the 12-bit images (pixel size 280 nm) thus obtained was segmented and reconstructed using Amira (Version 6.5, Thermo Fisher and Zuse Institute, Berlin, Germany).

### Focused ion beam-scanning electron microscopy

Serial focused ion beam-scanning electron microscopy (FIB-SEM) imaging was performed with a Zeiss Crossbeam 540 station (Carl Zeiss, Konstanz, Germany). Carbon-coated specimens were oriented inside the FIB-SEM chamber so that the viewing direction was aligned with the tendon longitudinal direction. A coarse cross-section was first milled with a 30 nA gallium beam at 30 kV acceleration voltage to provide a viewing channel for SEM observation. The exposed surface of this cross-section was fine-polished by lowering the ion beam current to 1.5 nA. Subsequently, the fine-polished block was serially milled by scanning the ion beam parallel to the surface of the cutting plane using an ion beam of 100 pA, 700 pA or 1.5 nA at 30 kV. After removal of each tissue slice, the freshly exposed surface was imaged at 1.5-3 kV acceleration voltage and 700 pA or 1 nA using both secondary electron (SE) and energy selective backscattered (EsB) detectors. The slice thickness was roughly equivalent to the lateral resolution of 2D images, ranging from 6-24 nm. In a fully automated procedure, the milling was combined with SEM imaging in sequence (imaging, then sectioning and reimaging) to collect thousands of serial images. Serial EsB images were aligned (image registration) with in-house python script in Anaconda (Austin, TX, USA). The same transformation matrix applied to each image of the EsB stack was then applied to the corresponding SE image stack. Following alignment, the images were digitally processed to increase contrast and reduce curtaining effects and noise levels. The resulting stacks of images were then segmented based on their structural features and reconstructed using Amira 3D (v 6.5; Thermo Fisher and Zuse Institute). Energy dispersive spectroscopy (EDS) analysis was performed using an Ultim Extreme silicon drift detector (Oxford Instruments, Abingdon, UK).

### Transmission electron microscopy

The same samples of tibialis cranialis tendons, prepared by either HPF or chemical methods and utilized for FIB-SEM, were subsequently examined by transmission electron microscopy. Tissues were stained in block with osmium tetroxide as noted previously and embedded in Spurr resin. Blocks were sliced (100 nm thickness) using a Leica Ultracut UCT ultramicrotome (Leica Microsystems) and sections were floated on a bath of anhydrous ethylene glycol (Electron Microscopy Sciences) to maintain the mineral phase of the tissue as optimally as possible. (48) Sections were collected on copper grids with a carbon support film. Images were obtained using a JEOL JEM-ARM200F electron microscope (JEOL, Ltd., Tokyo, Japan) operated at 200 kV, combining traditional TEM and scanning-TEM.

## Acknowledgements

The authors are grateful to Drs. Wolfgang Wagermaier and Richard Weinkamer for insightful discussions and to Birgit Schonert for technical support with sample polishing (all at the Max Planck Institute of Colloids and Interfaces, Golm, Germany).

## Supplementary Information

**Fig. S1:**
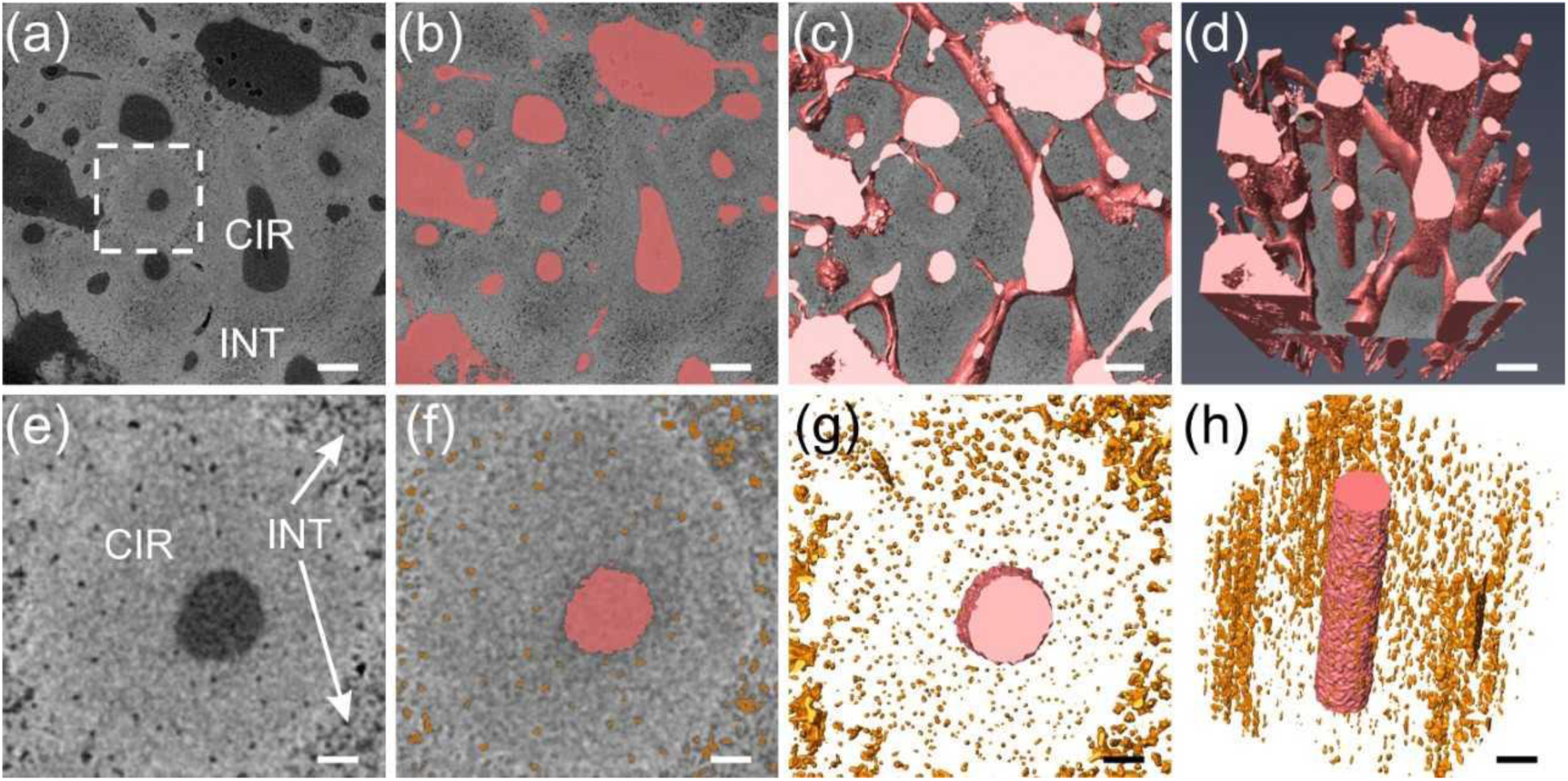
Micro-CT images of a distal tendon specimen prepared by chemical methods and stained with osmium tetroxide. **a**, A cross-sectional slice from 3D micro-CT data of the tendon with a voxel size of 1.2 µm^3^ shows a series of dark canals of various diameters traversing the tissue. The canals are surrounded by mineralized circumferential (CIR) tissue and such CIR tissue is itself separated by mineralized interstitial (INT) tissue. **b**, Unmineralized canals in **a** are highlighted in red. **c, d**, 3D surface rendering of unmineralized canals, presented at slightly different enlargements and angular aspects to demonstrate extensive interconnections or channels between canals. **e**, Enlarged image of the area marked in **a**, showing a single canal and numerous tenocyte lacunae. **f**, Unmineralized canal and lacunae in **e** highlighted in red and tangerine, respectively. **g, h**, 3D surface rendering of the canal (red) and lacunae (tangerine) in **f** viewed in transverse and approximately longitudinal profiles, respectively. Lacunae are elongated in longitudinal view and the lacunae long axes lie principally parallel to the canal. Both transverse and longitudinal views of the volume indicate lower density of lacunae within the circumferential zone compared to that within the interstitial region. Scale bars in panels **a**-**d**: 100 µm; in panels **e**-**h**: 25 µm.

**Fig. S2:**
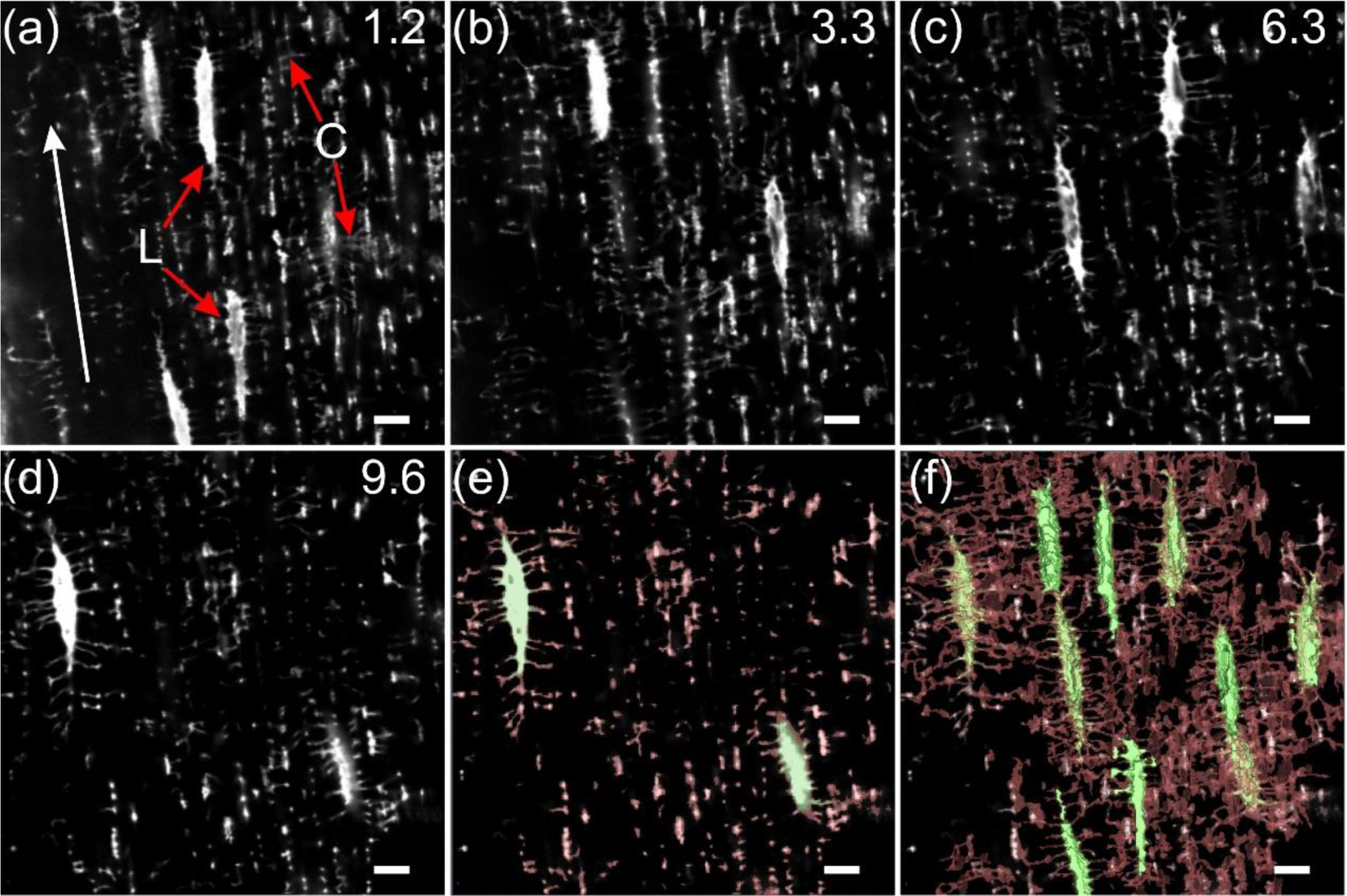
3D confocal images of a turkey distal tendon prepared by chemical fixation and osmium tetroxide staining. **a**-**d**, Representative slices of images from the 3D confocal volume show long, narrow tenocyte lacunae (L) and smaller canaliculi (C). The longitudinal direction of the tendon is marked by a white arrow. Slices are marked (upper right corner of an image) to designate the depth in microns from the tendon surface being viewed. **e**, Lacunae and canaliculi in **d** are highlighted in green and red, respectively. **f**, 3D surface rendering of the full volume of slices including **a**-**d** of lacunae and canaliculi demonstrates an extensive network resembling the lacuno-canalicular system in bone. Scale bar = 10 µm for all panels.

**Fig. S3:**
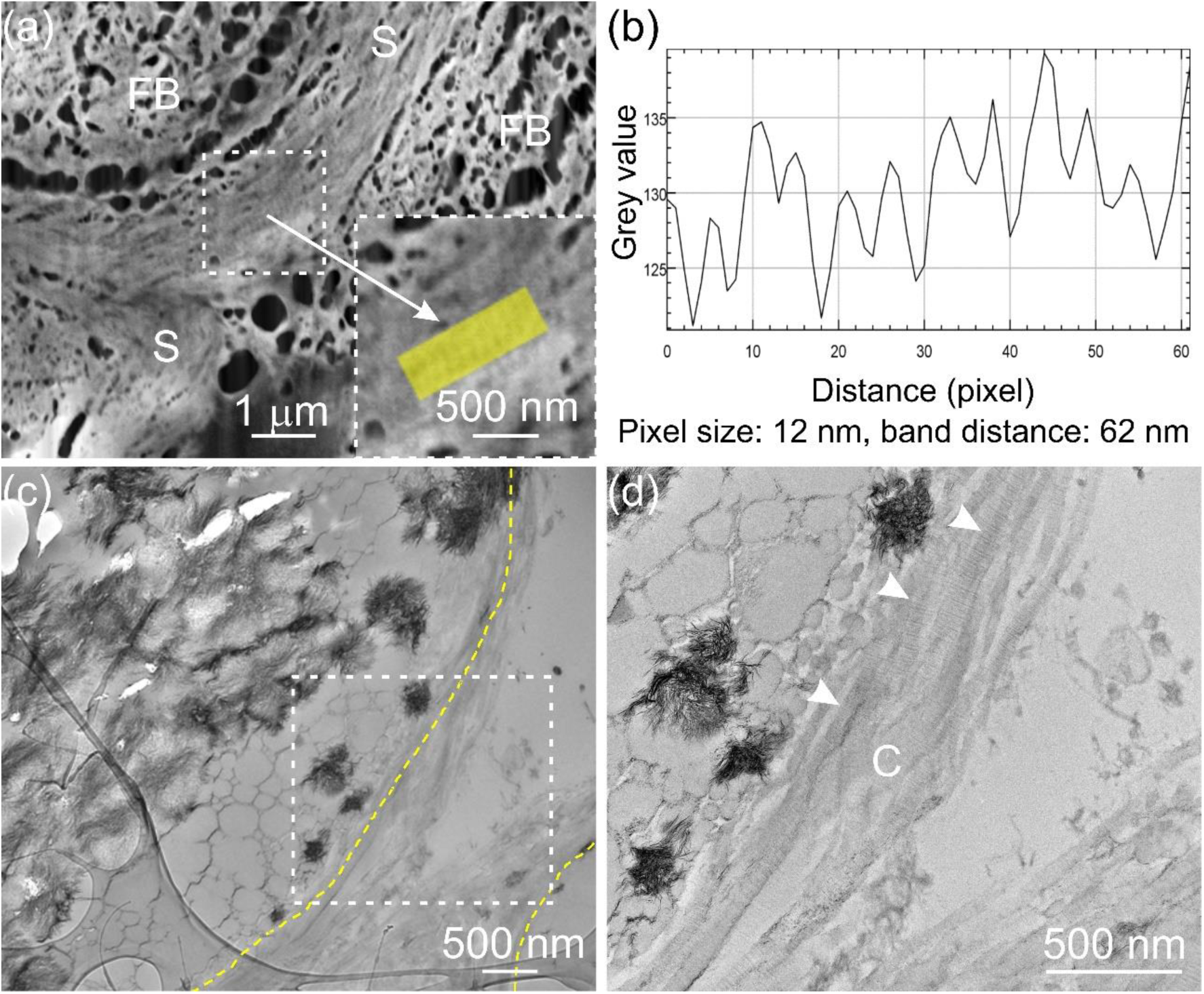
FIB-SEM and TEM images of the sheath region of a distal tendon specimen prepared by HPF and stained with osmium tetroxide-uranyl acetate. **a**, A transverse SEM image showing the sheath region (S) containing collagen fibrils oriented in the circumferential direction with respect to the long axes of several fibril bundles (FB). An area of the sheath (outlined by white dashes) was enlarged (arrow, lower right aspect of **a**) and a portion was examined (following a longitudinal midline through the yellow rectangular region) by Fiji to determine changes in image grey values. **b**, The characteristic periodic banding pattern of collagen fibrils (∼62 nm) was clearly observed on grey value analysis of this region. **c, d**, TEM images of a different region of the same tendon specimen shown in **a** illustrates the sheath region between two collagen fibril bundles. Yellow dashed lines denote the bundle boundaries. **d**, The enlarged TEM image of the region in **c** (white dashes) shows multiple collagen fibrils (C) with clear banding patterns (arrowheads) in the sheath region.

**Fig. S4:**
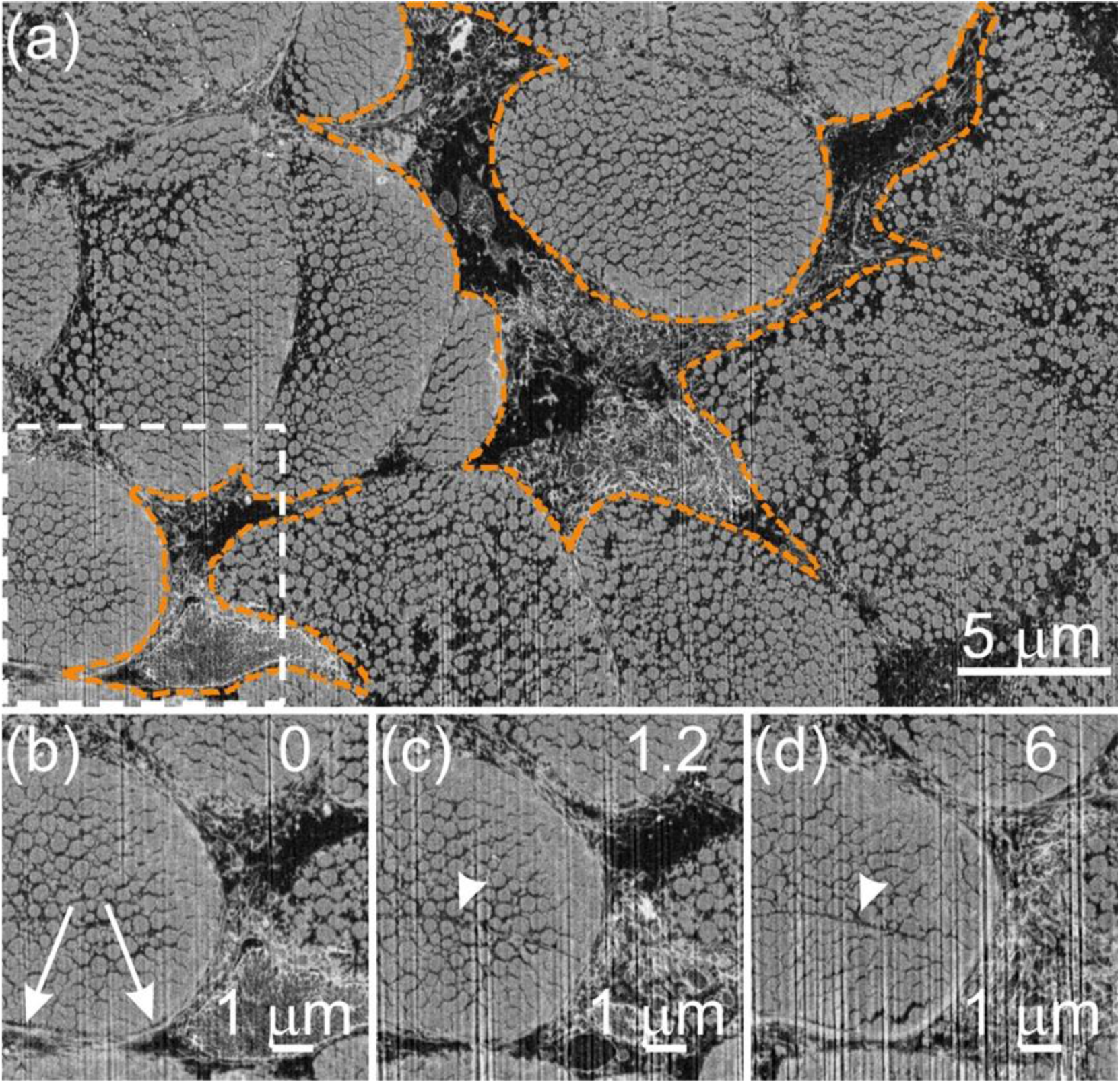
Cross-sectional FIB-SEM images of an unmineralized distal region of a tendon specimen prepared by HPF and stained with osmium tetroxide-uranyl acetate. **a**, An SE image showing ultrastructural aspects of two tenocytes (tangerine dashed lines) situated between multiple collagen fibril bundles. **b**, Enlarged SE image in **a** (white dashed frame) reveals possible cell processes (arrows) surrounding collagen fibril bundles. **c**,**d**, represent SE images obtained from the same volume of tissue as **b** but at different distances along the longitudinal direction of collagen fibril bundles. Numbers (top right corner of **c, d**) designate distance in µm from **b**. Putative canaliculi (arrowheads) were found occasionally traversing the collagen fibril bundles.

**Fig. S5:**
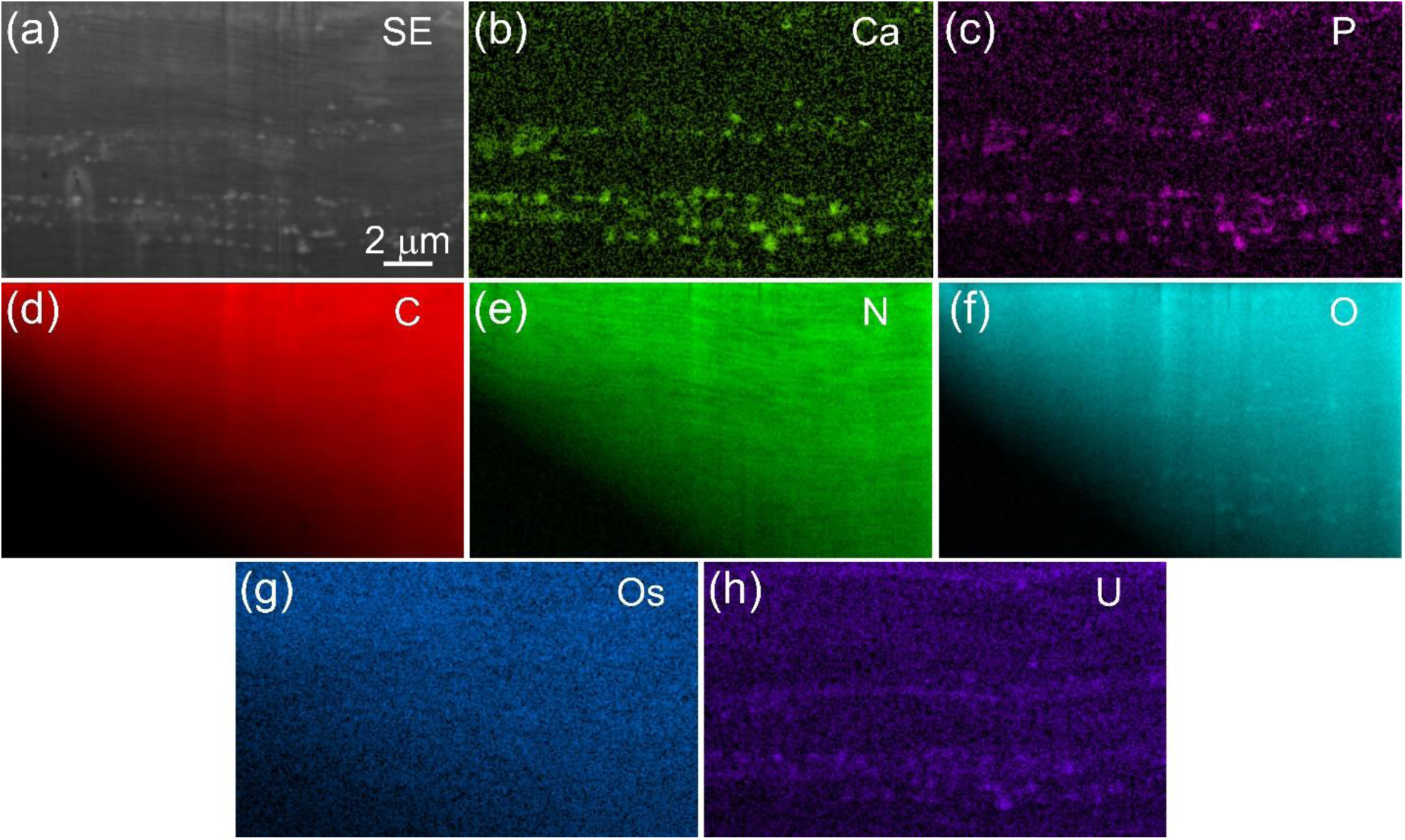
FIB-SEM image and corresponding EDS mapping at the mineralization front of a proximal tendon specimen prepared by HPF and stained with osmium tetroxide-uranyl acetate. **a**, An SE image of a longitudinal section of a tendon tissue showing mineral deposits (white) and collagen fibrils (grey). **b-h**, Corresponding EDS mapping images of different elements in **a**. The mineral regions are rich in calcium **b** and phosphate **c**. Nitrogen **e** shows slightly stronger signals in collagen while oxygen **f** has relatively higher intensity in mineral deposits. The signal intensities of carbon **d** and osmium **g** are homogeneously distributed across the tissue section. Uranyl **h** appears to have stronger association with mineral deposits.

## Movie legends

**Movie S1. 3D micro-CT imaging of a distal mineralized region of the tibialis cranialis tendon prepared by protocol 3.** The movie starts with cross-sectional images advancing from mineralized to unmineralized portions of the tissue. At 1 min, the movie changes to longitudinal images. Numerous dark black (empty) canals are observed with white (mineralized) circumferential and interstitial regions surrounding them. Dark black and grey areas about black are thought to be regions of the tissue without Spurr embedding resin or only limited in resin infiltration, respectively. The region also shows extensive channels that interconnect canals. Voxel resolution is 4.5 µm^3^.

**Movie S2. LSCM images viewed following the longitudinal direction of a distal tendon prepared by protocol 3.** Images progress from the tendon surface to deeper depths. In the movie with changing depth into the tissue, lacunae and canaliculi are visible and illustrate clearly the very extensive lacuno-canalicular network appearing similar to the osteocyte network in bone. Canaliculi are oriented principally perpendicular to the long axes of lacunae.

**Movie S3. FIB-SEM images (EsB) and 3D reconstruction of a small, fully mineralized distal tendon region in a transverse profile to the longitudinal direction of the tissue prepared by protocol 3.** Canaliculi originating from the lacunae of two representative tenocytes (orange) course through the ECM of the tendon composed here of mineralized (white) collagen fibril bundles. In 3D rendering, the canaliculi (tangerine) surround the bundles and clearly form circumferentially about them. Canaliculi from neighboring cells may interconnect.

**Movie S4. FIB-SEM images (SE) and 3D reconstruction of a small, fully mineralized distal tendon region in transverse profile to the longitudinal direction of the tissue prepared by protocol 3.** The movie shows numerous dark pores of various sizes and shapes throughout the ECM. These pores, reconstructed in 3D, comprise a vast system of secondary channels (cyan) interconnected and oriented in a perpendicular direction to canaliculi (tangerine) and directed parallel to mineralized (white) collagen fibrils and fibril bundles.

**Movie S5. FIB-SEM images (EsB) and 3D reconstruction of a small distal tendon region comprised of an interface between unmineralized and mineralized tissue zones.** The specimen is viewed in transverse profile to the longitudinal direction of the tissue prepared by protocol 1. The region is comprised of mineralized (white) and unmineralized (grey) collagen bundles together with a few intervening cells. 3D rendering shows tenocytes (tangerine) in linear disposition closely associated with mineralized collagen (blue) as the protein is secreted from the cells to form fibril bundles. The movie also suggests that there are many individual mineralized collagen fibrils in apparent units separated within a fibril bundle by dark thin structures yet to be identified; they may originate from the canaliculi but further work is required to establish their source.

**Movie S6. FIB-SEM images (SE) and 3D reconstruction of a distal tendon containing an interface between unmineralized and mineralized tissue zones.** The specimen is viewed in transverse profile to the longitudinal direction of the tissue prepared by protocol 2. The region consists of collagen fibril bundles, one more heavily mineralized (right half of image and in white) than the others (left half of image). Bundles are separated by a continuous dark, narrow space representing the sheath about the mineralized bundle. The more heavily mineralized bundle contains some fibrils yet to mineralize and which appear rather compact with indistinct borders. Fibrils comprising the less mineralized bundles are clearly separated from their neighbors by dark spaces which might be secondary channels. These same fibrils are not as compact as their counterpart unmineralized fibrils in the more heavily mineralized bundle. On 3D rendering of a portion of a less mineralized bundle, individual collagen fibrils (wine) of similar diameter, estimated to be ∼250 nm, and unfused with each other are found with mineral deposits (blue) both within (intra-) and outside (inter-fibrillar) the fibrils.

**Movie S7. FIB-SEM images (SE) and 3D reconstruction of a proximal tendon containing an interface between unmineralized and mineralized tissue zones.** The specimen is viewed in longitudinal profile following the long axis of the tissue prepared by protocol 2. The region is characterized by numerous collagen fibrils of various diameters and separated from one another by dark spaces, some of which contain mineral deposits (white and pseudo-colored red) of different sizes and shapes. The dark spaces are thought to represent the secondary channels that pervade the tendon and provide the space for fluid transport of mineral ion and mineral precursors. The mineral deposits in the interfibrillar collagen spaces are presumed to originate in association with putative matrix vesicles residing in secondary channels. Sites of intrafibrillar mineral deposition are marked by blue. In this movie and several others reconstructed from stacks of FIB-SEM images, intrafibrillar mineral sites are found in very close proximity to interfibrillar mineral sites, which are arranged linearly and in abundance near each intrafibrillar mineral deposit.

