## Supplemental figures for "Three-dimensional Structural Interrelations between Cells, Extracellular Matrix and Mineral in Vertebrate Mineralization"

This PDF file includes:

Figures S1 to S5 and legends

Movies S1 to S6 legends

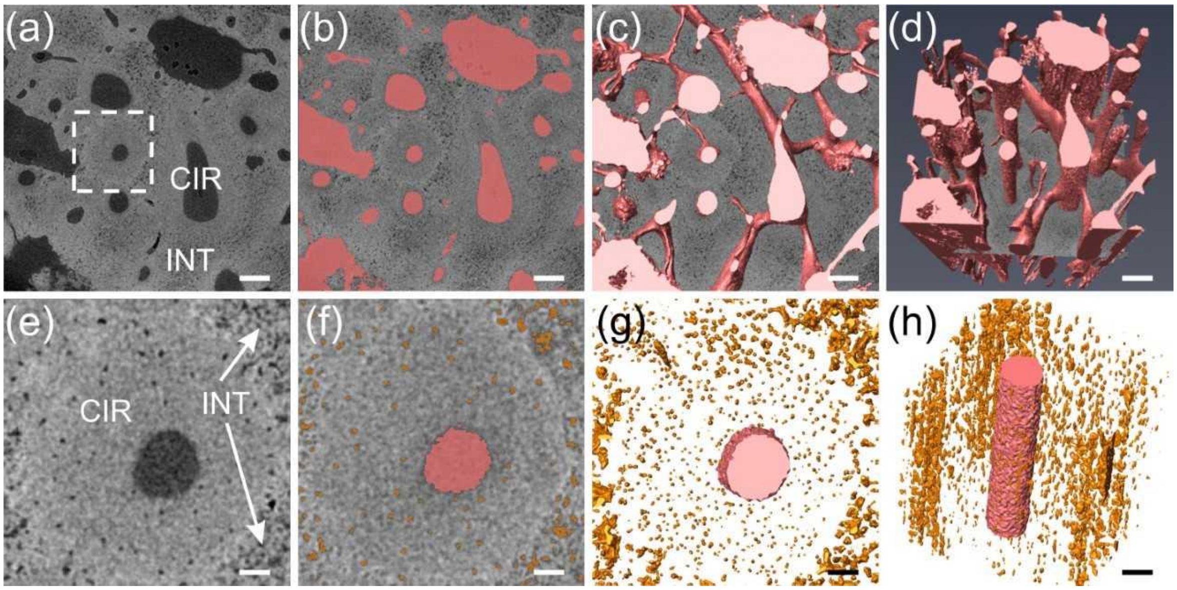


**Fig. S1: Micro-CT images of a distal tendon specimen prepared by chemical methods and stained with osmium tetroxide. a,** A cross-sectional slice from 3D micro-CT data of the tendon with a voxel size of 1.2 µm$^3$ shows a series of dark canals of various diameters traversing the tissue. The canals are surrounded by mineralized circumferential (CIR) tissue and such CIR tissue is itself separated by mineralized interstitial (INT) tissue. **b,** Unmineralized canals in **a** are highlighted in red. **c, d,** 3D surface rendering of unmineralized canals, presented at slightly different enlargements and angular aspects to demonstrate extensive interconnections or channels between canals. **e,** Enlarged image of the area marked in **a**, showing a single canal and numerous tenocyte lacunae. **f,** Unmineralized canal and lacunae in **e** highlighted in red and tangerine, respectively. **g, h,** 3D surface rendering of the canal (red) and lacunae (tangerine) in **f** viewed in transverse and approximately longitudinal profiles, respectively. Lacunae are elongated in longitudinal view and the lacunae long axes lie principally parallel to the canal. Both transverse and longitudinal views of the volume indicate lower density of lacunae within the circumferential zone compared to that within the interstitial region. Scale bars in panels **a**-**d**: 100 µm; in panels **e**-**h**: 25 µm.

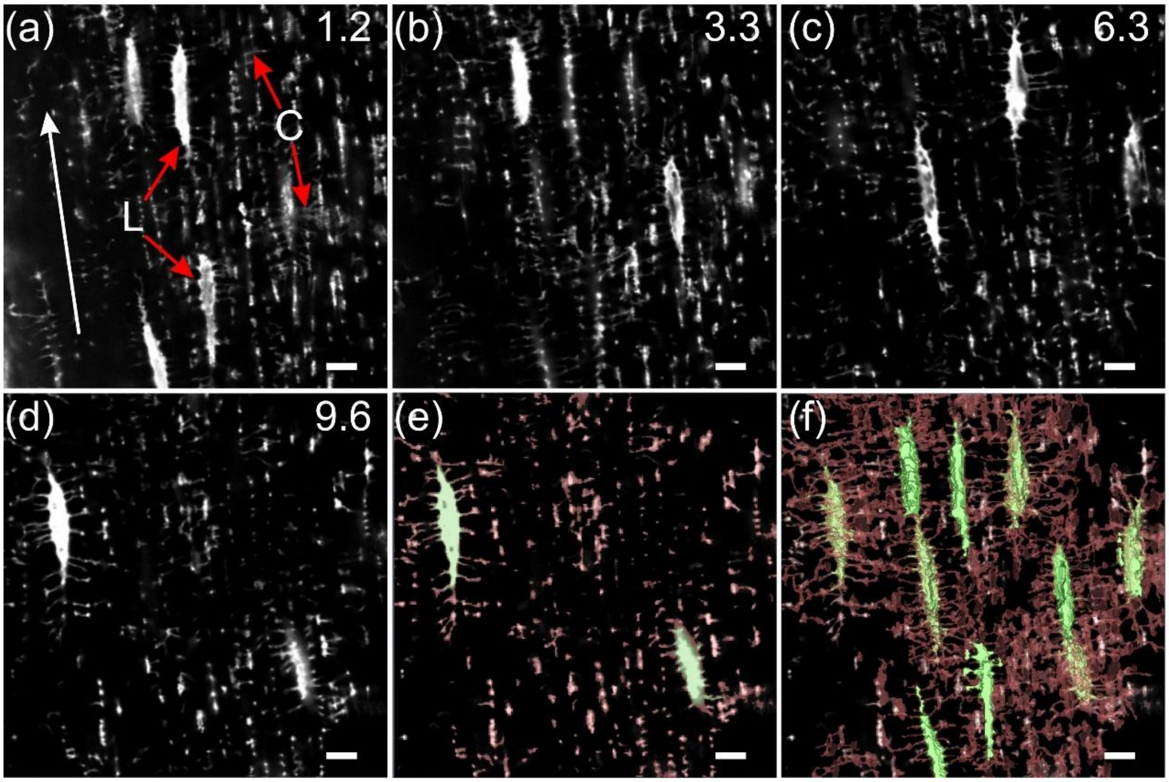


**Fig. S2: 3D confocal images of a turkey distal tendon prepared by chemical fixation and osmium tetroxide staining. a**-**d,** Representative slices of images from the 3D confocal volume show long, narrow tenocyte lacunae (L) and smaller canaliculi (C). The longitudinal direction of the tendon is marked by a white arrow. Slices are marked (upper right corner of an image) to designate the depth in microns from the tendon surface being viewed. **e,** Lacunae and canaliculi in **d** are highlighted in green and red, respectively. **f,** 3D surface rendering of the full volume of slices including **a**-**d** of lacunae and canaliculi demonstrates an extensive network resembling the lacuno-canalicular system in bone. Scale bar = 10 µm for all panels.

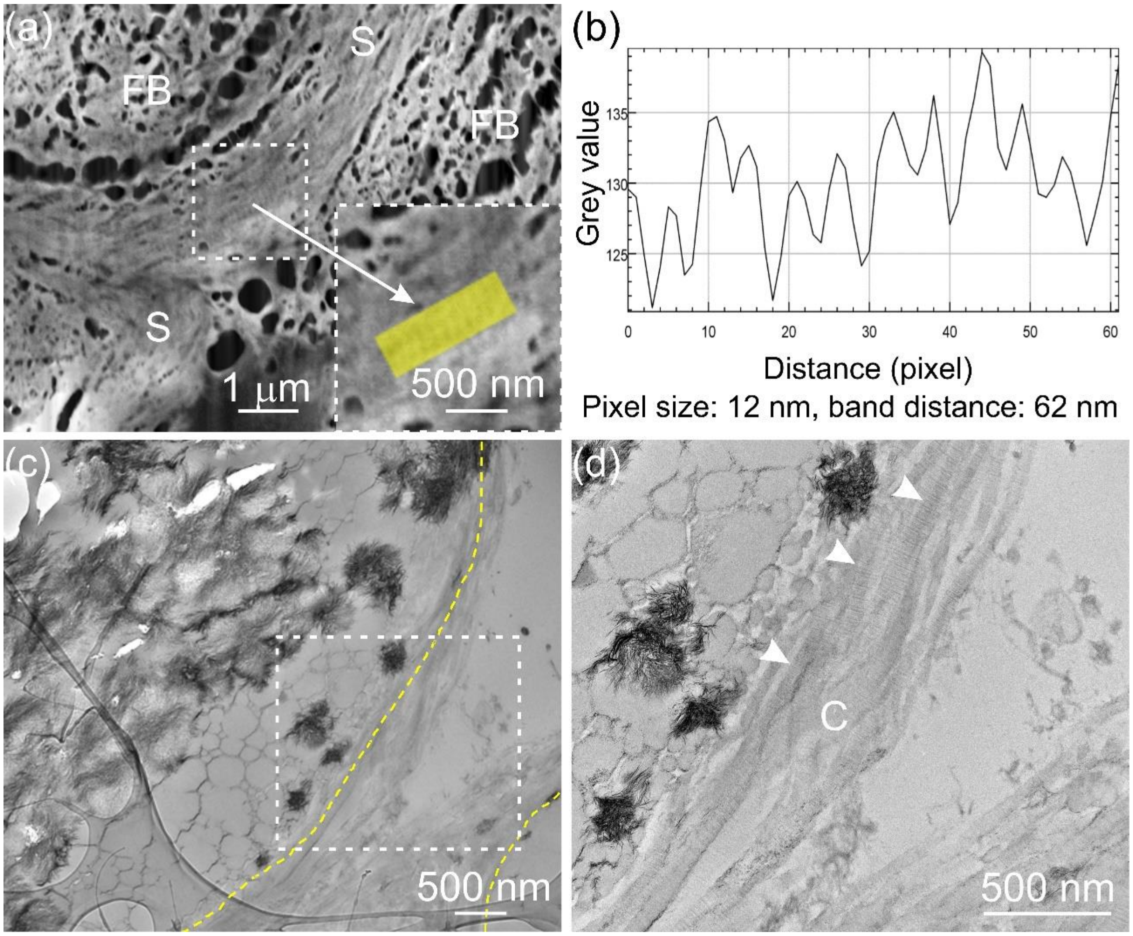


**Fig. S3: FIB-SEM and TEM images of the sheath region of a distal tendon specimen prepared by HPF and stained with osmium tetroxide-uranyl acetate. a,** A transverse SEM image showing the sheath region (S) containing collagen fibrils oriented in the circumferential direction with respect to the long axes of several fibril bundles (FB). An area of the sheath (outlined by white dashes) was enlarged (arrow, lower right aspect of **a**) and a portion was examined (following a longitudinal midline through the yellow rectangular region) by Fiji to determine changes in image grey values. **b,** The characteristic periodic banding pattern of collagen fibrils (~62 nm) was clearly observed on grey value analysis of this region. **c, d,** TEM images of a different region of the same tendon specimen shown in **a** illustrates the sheath region between two collagen fibril bundles. Yellow dashed lines denote the bundle boundaries. **d,** The enlarged TEM image of the region in **c** (white dashes) shows multiple collagen fibrils (C) with clear banding patterns (arrowheads) in the sheath region.

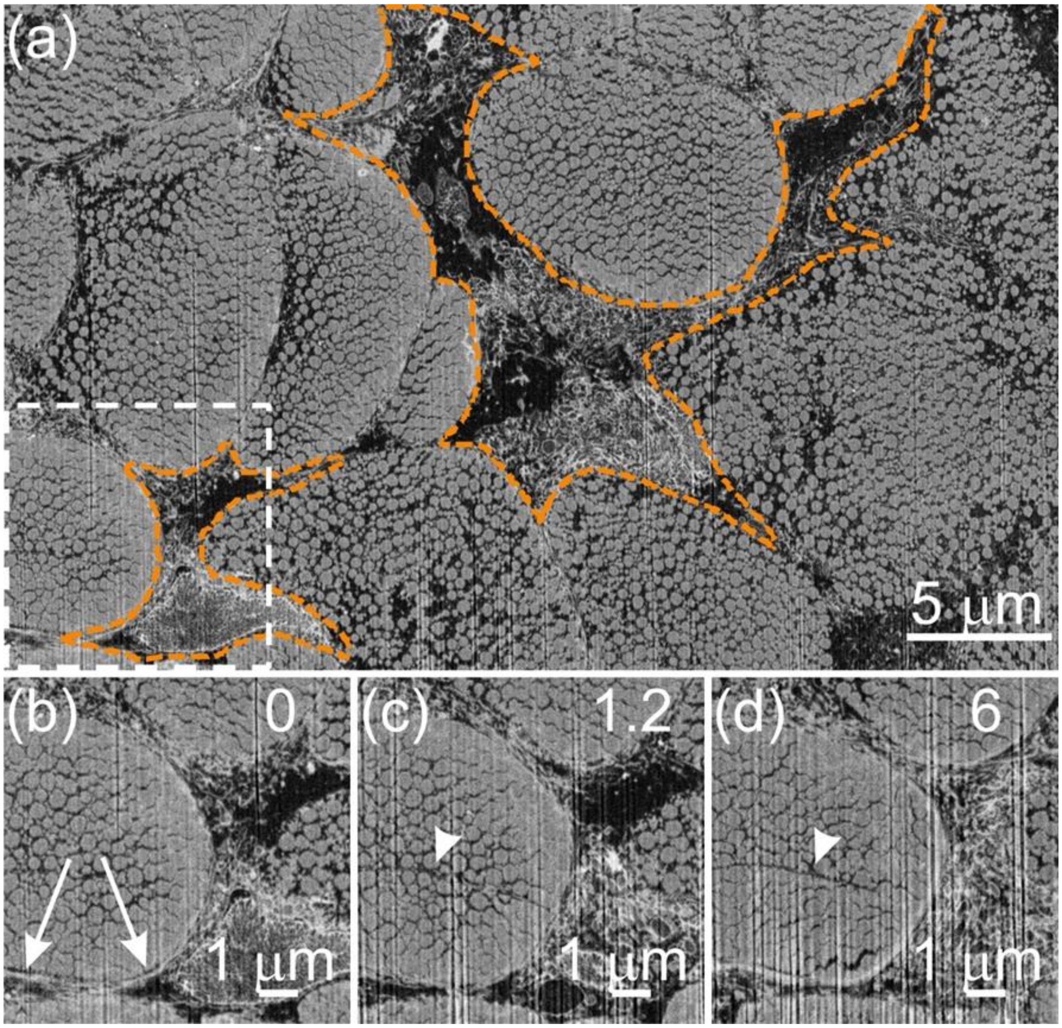


**Fig. S4: Cross-sectional FIB-SEM images of an unmineralized distal region of a tendon specimen prepared by HPF and stained with osmium tetroxide-uranyl acetate. a,** An SE image showing ultrastructural aspects of two tenocytes (tangerine dashed lines) situated between multiple collagen fibril bundles. **b,** Enlarged SE image in **a** (white dashed frame) reveals possible cell processes (arrows) surrounding collagen fibril bundles. **c**,**d,** represent SE images obtained from the same volume of tissue as **b** but at different distances along the longitudinal direction of collagen fibril bundles.

Numbers (top right corner of **c, d**) designate distance in µm from **b**. Putative canaliculi (arrowheads) were found occasionally traversing the collagen fibril bundles.

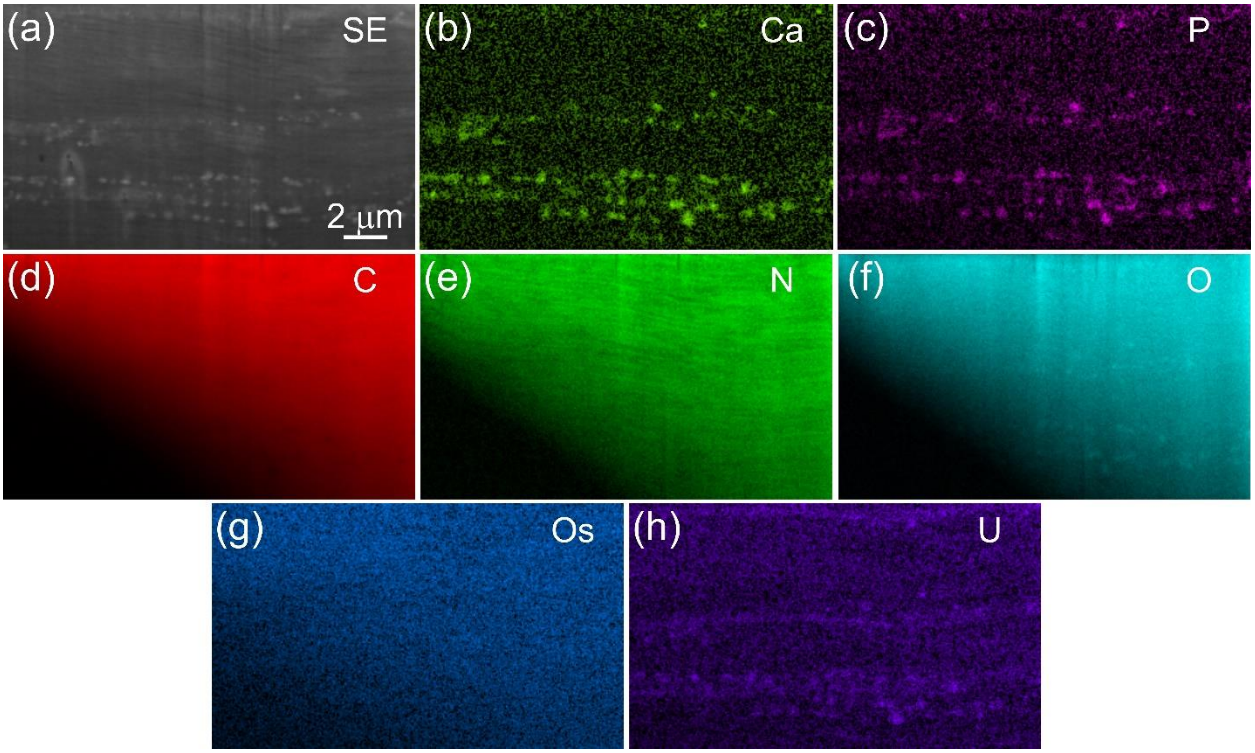


**Fig. S5: FIB-SEM image and corresponding EDS mapping at the mineralization front of a proximal tendon specimen prepared by HPF and stained with osmium tetroxide-uranyl acetate. a,** An SE image of a longitudinal section of a tendon tissue showing mineral deposits (white) and collagen fibrils (grey). **b-h,** Corresponding EDS mapping images of different elements in **a**. The mineral regions are rich in calcium **b** and phosphate **c**. Nitrogen **e** shows slightly stronger signals in collagen while oxygen **f** has relatively higher intensity in mineral deposits. The signal intensities of carbon **d** and osmium **g** are homogeneously distributed across the tissue section. Uranyl **h** appears to have stronger association with mineral deposits.
